# TaxSEA: an R package for rapid interpretation of differential abundance analysis output

**DOI:** 10.1101/2024.11.20.624438

**Authors:** Feargal J. Ryan

## Abstract

Microbial communities are essential regulators of ecosystem function, with their composition commonly assessed through DNA sequencing. Most current tools focus on detecting changes among individual taxa (e.g., species or genera), however in other omics fields, such as transcriptomics, enrichment analyses like Gene Set Enrichment Analysis (GSEA) are commonly used to uncover patterns not seen with individual features. Here, we introduce TaxSEA, an R package for taxon set enrichment analysis. TaxSEA integrates taxon sets from five public microbiota databases (BugSigDB, MiMeDB, GutMGene, mBodyMap, and GMRepoV2) to assess whether disease signatures, metabolite producers, or previously reported associations are enriched or depleted in a metagenomic dataset of interest. In-silico assessments show TaxSEA is accurate across a range of set sizes. When applied to differential abundance analysis output from Inflammatory Bowel Disease and Type 2 Diabetes metagenomic data, TaxSEA outperforms current tools and can rapidly identify changes in functional groups corresponding to known associations. We also show that TaxSEA is robust to the choice of differential abundance (DA) analysis package. In summary, TaxSEA enables researchers to efficiently contextualize their findings within the broader microbiome literature, facilitating rapid interpretation and advancing understanding of microbiome–host and environmental interactions.

## Background

Host-associated microbial communities, particularly the gut microbiota, play pivotal roles in immune regulation [1], drug metabolism [2] neural development [3], and nutrition [4] and are therefore frequently studied in medical research. The assessment of the gut microbiota commonly involves DNA sequencing, often with the aim of comparing the composition between samples. Comparing microbiota composition across samples is achieved through differential abundance (DA) analysis, which identifies taxa (e.g. species, genera, operational taxonomic units) which are present at varying levels between groups of samples (e.g. treatment vs control). However, interpreting alterations identified through DA analysis in humans is complicated by the fact that the gut microbiota is highly personalized yet functionally redundant [5]. For example, a reduction of species A in sample 1 and species B in sample 2 may have the same functional consequence (e.g. a decrease in the production of metabolite Xgy) but such changes appear unrelated when using taxonomic profiling alone. Current approaches to solve this problem, such as gutSMASH [6] and HUMAnN3 [7], profile gene clusters and pathways from metagenomic data and construct additional count matrices for further analysis. While useful, these approaches suffer from sparsity and high dimensionality issues in addition to requiring significant time and compute resources.

In the past, DA analysis was performed using either simple statistical tests (e.g. Wilcoxon rank sum test), or by adapting differential gene expression (DGE) approaches from transcriptomics (e.g. edgeR, DESeq2 [8]) although there are now multiple dedicated DA tools such as ALDEx2 [9], fastANCOM [10] and LinDA [11]. Results from DGE analyses are routinely further explored using gene set enrichment analyses (GSEA) which identify changes in sets of genes representing pathways (e.g. cell cycle, oxidative phosphorylation), disease associations or other functional groupings (e.g. genes regulated by the same transcription factor). This process was originally based upon the non-parametric Kolmogorov-Smirnov (KS) test which assesses whether the distribution of members in a predefined gene set differs from the overall dataset distribution [12]. GSEA takes seconds to run, requires minimal compute resources and allows for rapidly contextualizing findings and generating hypotheses from DGE analysis output. Currently, using this approach for DA analysis output demands specialized expertise, requiring users to create and format databases, manage taxonomic name mappings, and repurpose RNASeq tools which have been developed for other data types.

To address this gap, we have developed TaxSEA, an R package for Taxon Set Enrichment Analysis that directly utilizes output from common differential abundance analysis tools. TaxSEA is built on a GSEA-like framework, enabling the exploration of changes in user-defined taxon sets or pre-defined sets from five public microbiota databases, covering disease signatures, metabolite producers, and previously published associations. TaxSEA can query the NCBI Entrez API to convert between taxonomic naming schemas and is robust to the choice of differential abundance analysis software.

## Methods and implementation

TaxSEA is an open-source R package under the GNU GPL-3 licence. TaxSEA is based upon a database (TaxSEA-DB) constructed from the following sources:

- BugSigDB: a community-editable database of manually curated microbial signatures from published differential abundance studies[13]
- GMrepo v2: a curated human gut microbiome database with special focus on disease markers and cross-dataset comparison [14]
- gutMGene: a comprehensive database for target genes of gut microbes and microbial metabolites [15]
- mBodyMap: a curated database for microbes across human body and their associations with health and diseases [16]
- MiMeDB: the Human Microbial Metabolome Database [17]

For each source above, taxon sets were downloaded, taxonomic identifiers converted to NCBI taxonomic IDs by querying the NCBI Entrez API, made non-redundant and transformed into a named list of taxon sets in R where each element is a named set and the members NCBI taxonomic IDs. As MiMeDB does not provide API access or bulk downloads, a manually curated subset of taxon sets was downloaded. Similarly, only a subset of body sites from mBodyMap were included (oral cavity, skin and vaginal tract) [18]. Additionally, TaxSEA allows users to add custom sets to this database for testing. TaxSEA takes as input species or genus names together with a rank/value for each taxa (e.g. log2 fold change, correlation coefficient etc.). TaxSEA then utilizes the non-parametric Kolmogorov-Smirnov (KS) test to assesses whether the distribution of members in a predefined set differs from the overall distribution. Prior to testing TaxSEA filters the TaxSEA-DB to only include sets with a minimum number of taxa detectable in the input data (user settable, default=5). A P value is computed with the KS test for each taxon set and adjusted using the Benjamini-Hochberg procedure to control the False Discovery Rate (FDR). The output of TaxSEA is a table containing the name of the taxon set, the median rank for the detected members, the P-value, FDR, and members of that set which are detected in the data. This approach is adapted from the original Gene Set Enrichment Analysis (GSEA) from transcriptomics [12]

To assess TaxSEAs sensitivity, we used a signal implantation approach, a method shown to be to be effective for benchmarking differential abundance methods [19]. Specifically, we manipulated real fold change data generated metagenomic samples from the curatedMetagenomicData R package (v3.1) [20] by implanting a known signal and testing TaxSEAs ability to recover enriched sets. To accomplish this we generated a log2 fold change distribution by comparing randomly selected non-overlapping subsets of healthy adult samples (n=25 each) from the LifeLines-DEEP cohort (LLD, n=1,040) using the differential abundance software LinDA on default parameters from the MicrobiomeStat R package (v1.1) [11]. Each distribution was input to TaxSEA and any containing any pre-existing taxon set enrichment signatures (FDR < 0.05) were excluded from further analysis. This was repeated to create 1000 distributions representative of the null hypothesis in enrichment testing of DA-analysis output. Next, a distribution was randomly selected and modified by changing the values of the taxa in each set using the runif() function in R, which generates random values from a uniform distribution. The parameters for runif() were set according to the size of the taxon set, with minimum and maximum values defined by the effect size to be tested. The modified distribution was then used as input for TaxSEA to determine if it could detect the implanted signal. This procedure was repeated 1,000 times for each taxon set. R analyses were performed using R (v4.3.2). For the comparison between DA packages, fastANCOM (v0.0.4) and ALDEx2 (v1.34.0) both on default parameters were used. MSEA (v0.0.2) was performed in Python (v3.11.8). TSEA was run in the MicrobiomeAnalyst webtool using the microbiome-metabolite taxon sets (http://microbiomeanalyst.ca/MicrobiomeAnalyst). See Data Availability for code and underlying data.

## Results & Discussion

TaxSEA is an R package to identify enrichments or depletions of sets of taxa (e.g. species, genera) corresponding to functional groups (e.g. producers of a particular metabolite) (**Fig. 1**). TaxSEA is built upon publicly available databases which we have collated together into the TaxSEA database (TaxSEA-DB, **Fig. 1A**). In total this database contains 2,529 taxon sets made up of 2,785 individual taxa from GMRepoV2 [14], GutMGene [15], mBodyMap [16], MiMeDB [17] and BugSigDB [13]. During the development of TaxSEA, we found that public databases disproportionately favour the human gut microbiota. Therefore, our focus here is on evaluating TaxSEA to interpret differential abundance (DA) analysis in that context. Nonetheless, TaxSEA can in principle handle samples from any environment, and offers users the flexibility to add custom taxon sets for testing. To aid with interpretation and usability we have broadly divided taxon sets from these databases into three categories for reporting enrichment signatures – health and disease signatures (GMRepoV2, mBodyMap), metabolite producers (MiMeDB, GutMGene) and previously published microbiota associations (BugSigDB). Select taxon sets from mBodyMap covering non-gut sites (mouth, skin, vaginal tract) were included as the presence of non-gut associated bacteria can be a feature of dysbiosis [21, 22].

**Figure 1:**
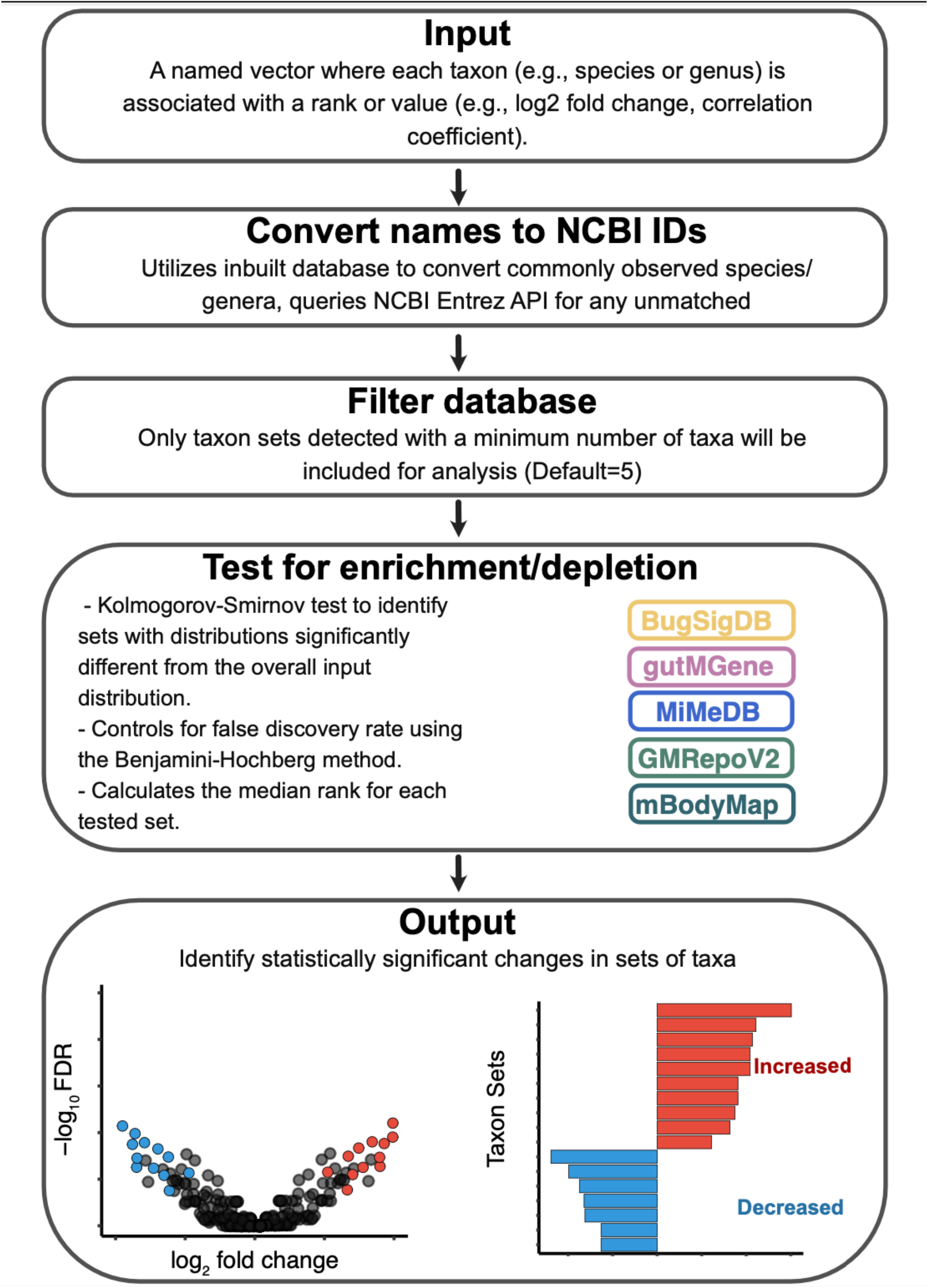
Overview of the taxon set enrichment analysis workflow using TaxSEA.

We first sought to evaluate whether TaxSEA is accurate. While many metagenomic analysis tools have been benchmarked with efforts like the Critical Assessment of Metagenome Interpretation (CAMI) [23], enrichment analysis present additional challenges. A single bacterial species will produce many metabolites, often as part of related metabolic processes, and certain species develop cross-feeding relationships and so may form a co-occurring group. Developing a ground truth dataset in this context is challenging as such overlaps represent biological realities but complicate the calculation of false negatives from enrichment testing. As such we assessed the accuracy of TaxSEA by first evaluating sensitivity (i.e., true positive rate, TPR) with an in-silico approach, and second by testing the ability of TaxSEA to recover known and biological plausible signals in real data.

TPR was evaluated by implanting an enrichment signal in real data from the LifeLines-DEEP dataset (LLD, n=1,040) of healthy adult gut metagenomic samples. For example implanting a set with a median fold-change of 1.5x (**Fig. 2A**) or 2.5x (**Fig. 2B**). The TPR of TaxSEA improves with higher fold changes and larger set sizes, reflecting the algorithm’s ability to capture more prominent biological signals effectively (**Fig. 2C**). For taxon sets with fewer than 10 members, the TPR was 11.9% at a 1.5x fold change but increased to 85.8% at 3x fold change. In medium-sized taxon sets (10-50 members) the performance improves substantially, with a TPR of 47.01% at 1.5x, 86.9% at 2x and >97% beyond 2.5x (**Fig. 2C**). Interestingly, larger taxon sets (more than 50 members) show a poorer performance than medium sets with TPR of 29.7% at a 1.5x change and 72.3% at 2x potentially reflecting a dilution of signal as the size of the set. Overall, this signal implantation evaluation found that TaxSEA demonstrates high sensitivity when taxon sets have a greater than 2.5x change, particularly in sets larger than 10 members.

**Figure 2:**
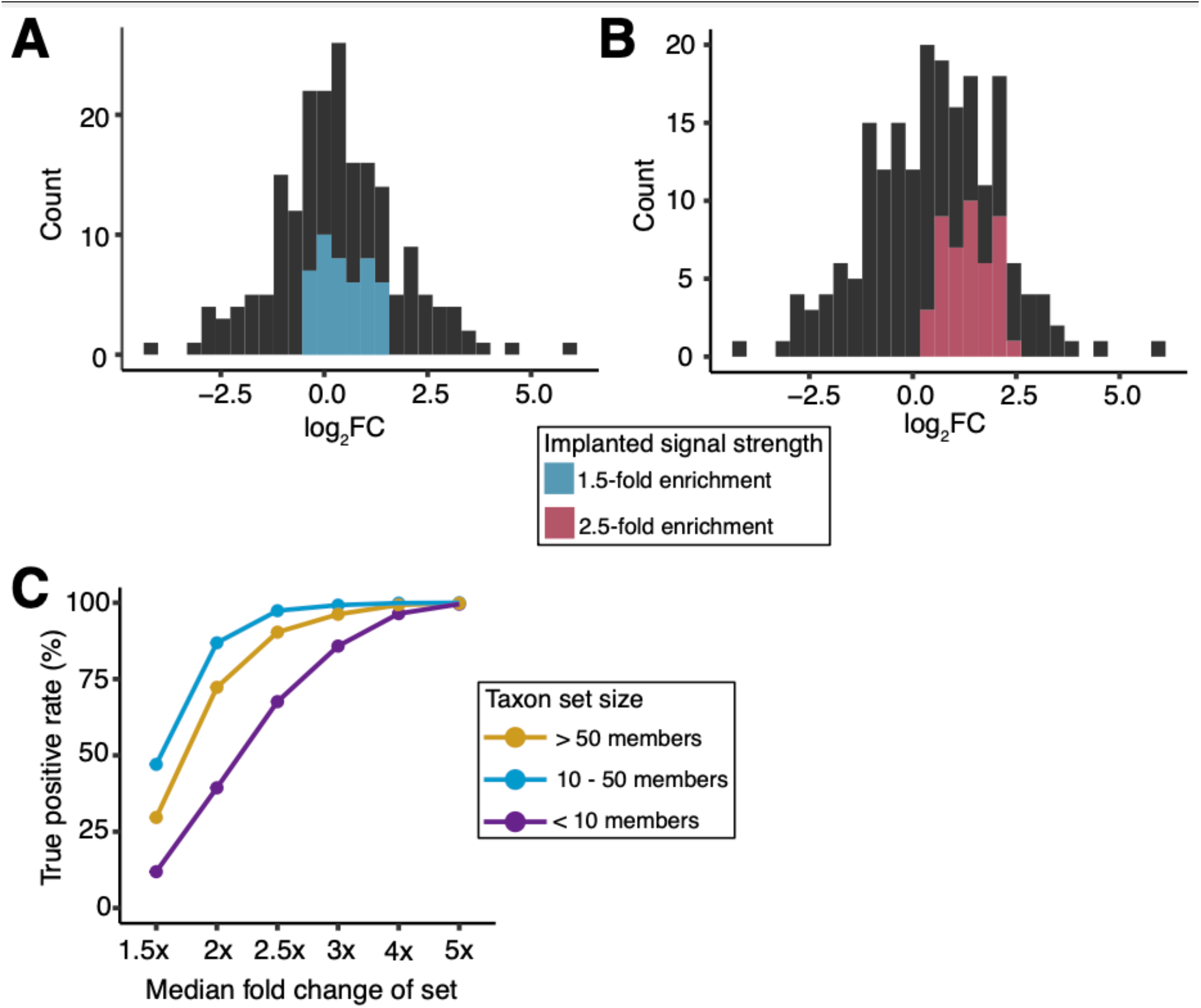
Representative examples from in-silico signal implantation assessments, displaying fold change distributions with (A) a 1.5x increase and (B) a 2.5x increase for a taxon set. (C) True positive rate (TPR) of TaxSEA at different fold changes and taxon set sizes.

Next, we sought to evaluate whether TaxSEA could detect biologically plausible associations in real data from the curatedMetagenomicData R package [20]. To avoid issues with overlapping testing and training data we excluded disease signatures from this evaluation as GMRepoV2 is built from public metagenomic datasets. First, we re-analysed data from the second phase of the Human Microbiome Project (Lloyd-Price *et al*. [24]) which compared the faecal metagenome of Inflammatory Bowel Disease (IBD) patients (n=99) and controls (n=27). DA-analysis with LinDA identified 5 species which were significantly decreased among IBD patients compared to controls (FDR < 0.1, **Table S1**, *Roseburia hominis, Ruminococcus torques, Ruminococcus bromii, Firmicutes bacterium CAG 83, Roseburia sp. CAG 18*2). TaxSEA detected significant (FDR < 0.1, median absolute fold change > 2.5x) alterations in 6 taxon sets marked by a depletion in producers of short chain fatty acids (SCFA) such as butyrate, acetate, and propionate driven by decreases in taxa such as *Faecalibacterium prausnitzii*, multiple *Roseburia* spp., *Prevotella copri*, among others (**Table S2, Fig 3A**). A depletion of SCFA producers is hallmark of the IBD microbiota [25]. Notably, several of these SCFA-producing species did not meet the threshold for being statistically significant alone (**Table S1, Fig 3B**) reflecting a significant shift in functional capacity shared across different species not observable in taxonomic composition. Next, we applied TaxSEA to the Qin *et al*. data which profiled the faecal metagenome of Type 2 Diabetes (T2D) patients (n=170) and controls (n=174) [26]. Here, DA-analysis identified 40 species which were significantly different between T2D and controls (FDR < 0.1, 26/40 increased in T2D, **Table S3**). Using the per-species fold changes as input, TaxSEA detects a significant depletion of producers of 2 metabolites (FDR < 0.05, median absolute fold change > 2.5x) - indole-3 propionic acid (IPA, **Fig. 3C**), and phenylalanine. (**Table S4**). IPA is a microbially produced tryptophan metabolite which directly modulates insulin secretion and is being investigated as biomarker for T2D [27, 28]. Finally, we sought to evaluate whether the choice of DA analysis package to generate the input ranks for TaxSEA would impact detection of these signatures. After reperforming each of the above DA-analyses with ALDEx2 and fastANCOM, we found that the resulting TaxSEA P values for each set were highly correlated (Pearson Cor > 0.97, **Figure 3D-E**) and that the number and identity of the taxon sets detected as significantly altered was identical across the 3 DA packages. Thus, TaxSEA can detect biological relevant alterations in gut metagenomic data and is robust to the choice of DA analysis method.

**Figure 3:**
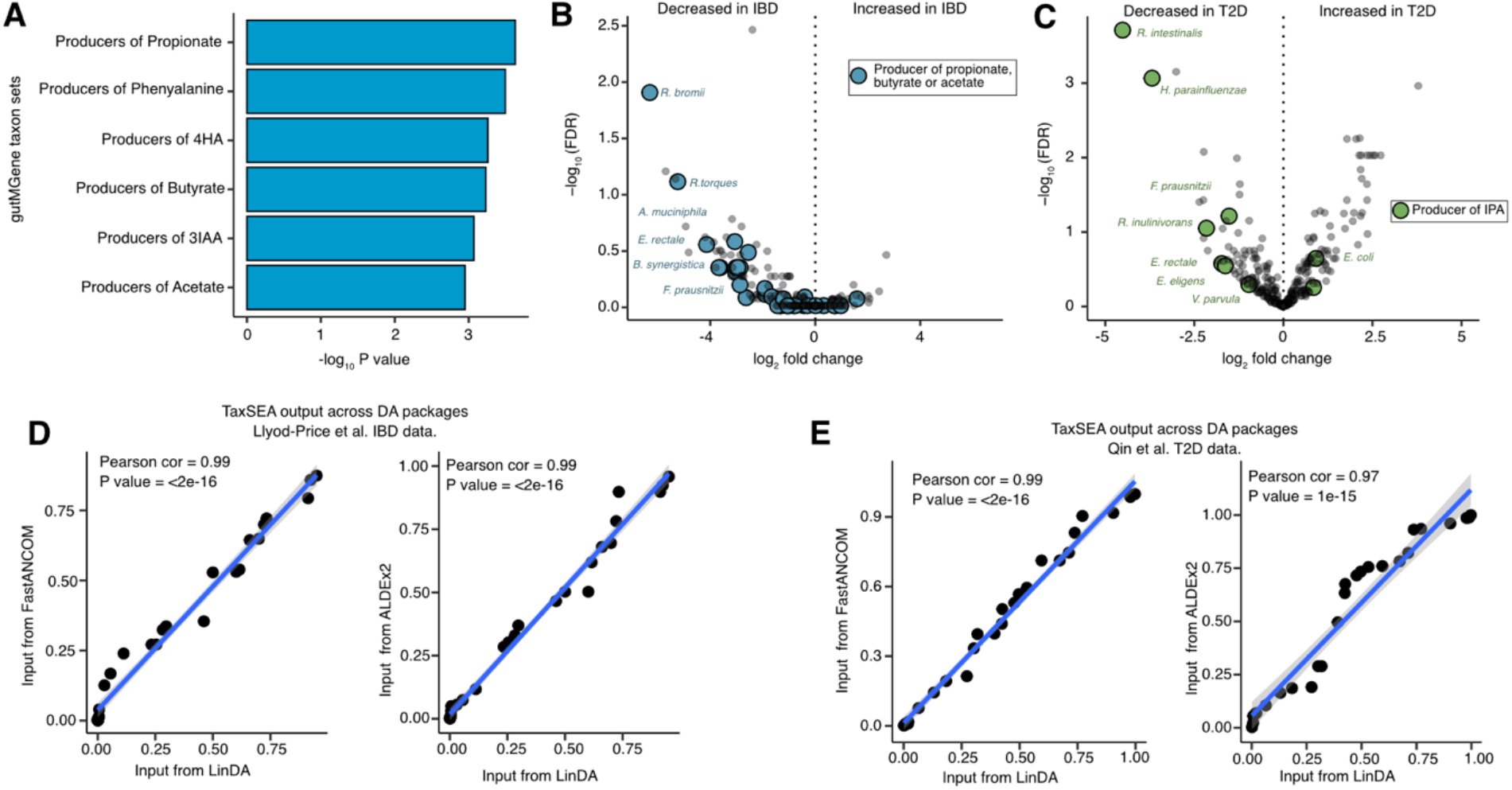
(A) Barplot of statistically significantly depleted taxon sets (FDR < 0.1) in inflammatory bowel disease (IBD) compared to controls in the Lloyd-Price et al. dataset. Abbreviations: 4HA = 4-Hydroxyphenylacetic acid; 3IAA = Indole-3-acetic acid. (B) Volcano plot showing depletion of short-chain fatty acid (SCFA) producers (i.e., propionate, butyrate, or acetate producer taxon sets) from the Lloyd-Price et al. dataset. (C) Volcano plot highlighting the depletion of producers of Indole-3-propionic acid (IPA) in type 2 diabetes (T2D) patients from the Qin et al. dataset. (D) Scatter plots showing correlations between p-values from TaxSEA across three differential abundance analysis tools (LinDA, FastANCOM, and ALDEx2) using (D) Lloyd-Price et al. dataset and (E) Qin et al. dataset.

We identified two methods in the literature that perform comparable enrichment testing on microbiota data. However, both rely on a different statistical approach, using an over-representation analysis (ORA) framework to assess the overlap between input species and predefined taxon sets; the Taxon Set Enrichment Analysis (TSEA) module in MicrobiomeAnalyst pipeline [29, 30] and the microbe-set enrichment analysis (MSEA) tool [31]. MSEA calls upon a combination of taxon sets defined by literature mining together with the Disbiome database [32] whereas TSEA incorporate sets from a variety of sources, including some of the same as the TaxSEA-DB [29, 30]. Although in both tools the user is limited in the ability of the user to test custom sets or other public databases. Despite these methodological differences, we sought to evaluate how these tools compared to TaxSEA. For each the IBD and T2D DA comparisons above, we extracted the list of DA species (FDR <0.1), split by fold change into increased and decreased species and then used each as input. Neither MSEA nor TSEA detected any significantly (FDR < 0.1) enriched/depleted taxon sets. As noted above, the depletion of butyrate producers we detected was in part driven by taxa which did not meet a threshold for statistical significance alone highlighting the limitation of requiring a strict cut-off as part of an ORA test. Nonetheless, an ORA based approach ORA may be more suitable when taxa are analysed without associated ranks. Ngyuen and colleagues previously reported on a technique known as Competitive Balances for Taxonomic Enrichment Analysis (CBEA) [33] which offers the ability to by generate enrichment scores for taxon sets in individual samples. When combined with TaxSEA and that merged database we present here, this allows researchers to identify group-level differences in taxon sets, which can then be examined at the sample level for deeper biological insights.

## Conclusion

Here, we present TaxSEA, an R package that enables researchers to rapidly interpret and contextualize DA analysis results. While our focus has primarily been on the gut microbiota, TaxSEA is versatile and can be applied to any microbial community, including non-human-associated environments. This work demonstrates that TaxSEA successfully extracts implanted signals and recovers known associations from DA outputs. The primary limitation of TaxSEA is that it is inherently reliant on the quality and comprehensiveness of the underlying databases. As microbiota research advances and databases grow larger, we expect enrichment approaches will similarly grow in their utility. We encourage users view ORA, KS-based enrichment, and sample-specific approaches like CBEA as complementary tools that together provide a robust framework for analysing microbiota associations and identifying shifts in functional capacity.

## Availability of data and materials

TaxSEA is open source and available as an R package on GitHub alongside all code and data for generating the results presented here (https://github.com/feargalr/TaxSEA). Benchmarking and evaluation code can be found on GitHub (https://github.com/feargalr/TaxSEA_benchmarking).

## Contributions

TaxSEA was conceived, implemented, and evaluated by FJR, who also authored the manuscript.

## Competing interests

The authors declare that they have no competing interests.

## Funding

This study was financially supported by grants from the National Health and Medical Research Council of Australia (APP2017404).

## Acknowledgements

The authors would like to thank David Lynn, Calum Walsh, Sam Forster, and Steven Taylor for their insightful discussions and valuable feedback throughout the development of this work. Large language models (LLMs) were utilized to assist with grammar and spelling corrections, as well as for code documentation.

---

**Table S1:** Differential abundance analysis output (LinDA) for the Lloyd-Price et al. dataset, comparing type 2 diabetes (T2D) samples to controls.

**Table S2:** TaxSEA output for the Lloyd-Price et al. dataset, comparing IBD samples to controls.

**Table S3:** Differential abundance analysis output (LinDA) for the Qin et al. dataset, comparing T2D samples to controls.

**Table S4:** TaxSEA output for the Qin et al. dataset, comparing T2D samples to controls.

## Notes

https://github.com/feargalr/TaxSEA

### Competing Interest Statement

The authors have declared no competing interest.

